# Dynamical analysis of a model of BCL-2-dependent cellular decision making

**DOI:** 10.1101/2025.07.08.663634

**Authors:** Ielyaas Cloete, Tomás Alarcón

## Abstract

The BCL-2 protein family governs critical cell-fate decisions between survival, senescence, and apoptosis, yet the dynamical principles underlying these choices remain poorly understood. Here, we integrate mathematical modeling, bifurcation analysis, and stochastic simulations to dissect how BCL-2 network architecture encodes multistability and fate plasticity. Our coarse-grained model reveals tristable regimes requiring cooperative BH3-only and anti-apoptotic BCL-2 interactions, with stochastic fluctuations driving heterogeneous fate commitments in genetically identical cells. Comparative analysis of mechanistic models demonstrates that while bistability emerges from canonical BCL-2 interactions, robust tristability requires additional regulatory constraint, explaining the metastability of senescence in stress responses. Hybrid models further show that BH3-only binding cooperativity enables multistability, but physiological senescence likely depends on additional control mechanisms. These results establish a unified framework linking molecular interactions to cell-fate dynamics, with implications for targeting apoptosis resistance in disease.

## Introduction

Apoptosis, or programmed cell death, is a fundamental process in development that regulates tissue homeostasis, eradicates autoreactive immune cells, and eliminates damaged and infected cells Kerr et al. (1972); Elmore (2007). Cellular senescence (hereafter referred to as senescence), a state of permanent cell cycle arrest, is a normal process of aging and may be transiently induced during development and tissue remodeling Hayflick (1965); Muñoz-Espín et al. (2013); Basu (2022). Dysregulated apoptosis and senescence can lead to pathological diseases and are often implicated in age-related diseases, including cancer, type-2 mellitus diabetes, neurodegenerative disorders, and cardiovascular diseases Childs et al. (2014); Cerella et al. (2016); Wanner et al. (2020). Apoptosis and senescence are induced by extrinsic and intrinsic stimuli, including the activation of death receptors and various external and internal stressors, such as DNA damage, oxidative stress, and telomere shortening Fulda and Debatin (2006); Gude et al. (2018).

Although apoptosis and senescence are processes that lead to distinct cell fates, their functional and mechanistic regulation is closely related. In certain cases, cells undergo apoptosis in response to high levels of stress, while senescence appears to be a consequence of moderate levels of stress Chen et al. (2000); Song et al. (2005); Vousden and Lane (2007); Spallarossa et al. (2009); Childs et al. (2014); Victorelli et al. (2023). Therapeutic interventions, including chemotherapeutics and ionizing radiation, have also been shown to induce a senescence phenotype, characterized by resistance to apoptosis Saleh et al. (2020). This suggests that the signaling pathways that regulate apoptosis and senescence are tightly interconnected. Indeed, the BCL-2 family of proteins, prominent regulators of mitochondrial-dependent apoptosis, also regulate senescence Roger et al. (2021); Basu (2022).

Based on protein structure and function, BCL-2 proteins are partitioned into pro-apoptotic BCL-2 proteins (BAX, BAK, etc.), anti-apoptotic BCL-2 proteins (BCL-2, BCL-xL, MCL-1, etc.), and sensitizer/activator BH3-only BCL-2 proteins (BID, BIM, PUMA, NOXA, etc.) Hata et al. (2015). The induction of apoptosis through the intrinsic pathway triggers the oligomerization of pro-apoptotic BAX and BAK, resulting in mitochondrial outer membrane permeabilization (MOMP) Kalkavan and Green (2018). Under normal cellular conditions, anti-apoptotic BCL-2 proteins sequester pro-apoptotic BAX and BAK, thereby preventing MOMP and preserving mitochondrial integrity. Intrinsic apoptotic signals activate BH3-only proteins, which neutralize antiapoptotic BCL-2 proteins, liberating pro-apoptotic mediators to trigger MOMP Delbridge et al. (2016); Radha and Raghavan (2017); Adams and Cory (2018); Singh et al. (2019). A subset of BH3-only proteins may directly activate pro-apoptotic BAX and BAK, though this mechanism remains controversial Luo et al. (2020).

The seminal work by Wang Wang (1995) first demonstrated that human fibroblasts resist apoptosis despite decreased levels of BCL-2 in younger cells, suggesting that anti-apoptotic BCL-2 proteins promote senescent cell survival. Subsequent studies revealed that oxidative stress transiently upregulates BCL-2 during growth arrest but not apoptosis, while BCL-2 overexpression induces premature senescence in murine fibroblasts and accelerates Ras-induced senescence in human fibroblasts without affecting cell cycle arrest Bladier et al. (1997); Tombor et al. (2003); López-Diazguerrero et al. (2006). Taken together, these findings establish a direct mechanistic link between BCL-2 upregulation and cellular senescence.

BCL-2 proteins play a crucial role in regulating the cellular fate decision between apoptosis and senescence. Senescent cells are often regarded as apoptosis-resistant, and BCL-2 protein activity has been implicated in cell fate decisions across diverse cell types Childs et al. (2014). Upregulation of anti-apoptotic BCL-2 family members is a consistent feature of senescence in multiple cell types Spaulding et al. (1999); Yosef et al. (2016); Zhu et al. (2016, 2017); Baar et al. (2017); He et al. (2020); Hu et al. (2022). Furthermore, chemo-and radiotherapy can induce early-onset senescence, known as therapy-induced senescence, which may increase tumor growth and invasion, potentially leading to treatment resistance Schmitt et al. (2022); Wang et al. (2022). Since upregulation of anti-apoptotic BCL-2 proteins is commonly associated with many cancers, senescence, and multiple age-related diseases, numerous studies have focused on anti-apoptotic BCL-2 proteins for cancer therapy and senotherapy.

The mechanisms governing the choice between apoptosis and senescence are not fully understood, including the regulatory role of BCL-2 proteins in these cell fate decisions. Mathematical and computational models are essential for interpreting complex molecular signaling dynamics and elucidating how these signaling networks drive cell fate decisions. Numerous computational models have studied the BCL-2-dependent apoptotic signaling network, including the cellular decision between cell survival and cell death Fussenegger et al. (2000); Eissing et al. (2004); Bagci et al. (2006); Legewie et al. (2006); Chen et al. (2007); Cui et al. (2008); Song et al. (2019); Cloete et al. (2023). Aspects of the intrinsic senescent signaling network have also been encoded in computational models Chong et al. (2018); Kim et al. (2019). Furthermore, a telomere-dependent binding model has also been developed to explain the apoptosis-senescence cell fate choice Arkus (2005). However, these models have not included the BCL-2 signaling network and are only sufficient to describe systems with two cellular states, cell survival or cell death (apoptosis). Thus, a new theoretical study of BCL-2-dependent apoptosis-senescence cell fate decision, including a study of the underlying dynamics that govern these distinct cell fates, is timely.

Here, we present a novel BCL-2-dependent model of cell fate decision making, including detailed analyses of the underlying dynamical properties of the system. Specifically, we examine how interactions between BCL-2 family proteins contribute to the robustness of cellular decision-making. First, we perform parametric perturbations to analyze how the stability structure changes with key parameters, identifying critical thresholds and the mechanisms driving multistability. Next, we perform an extensive exploration of the phase space to analyze how key dynamical structures confer robustness to the various cell fates. We then perform stochastic simulations to model the intrinsic noise associated with BCL-2 protein interaction, enabling the quantification of robustness and phenotypic stability Skommer et al. (2010). Finally, we map our coarse-grained model to high-dimensional mechanistic models of BCL-2 signaling, increasing the biological interpretability of our model.

## Methods

### Model

Conceptual models of BCL-2 protein interaction have been presented previously Chipuk and Green (2008); Leber et al. (2010); Llambi et al. (2011); Hata et al. (2015), see Shamas-Din et al. (2013) for a review. The ‘Embedded together’ framework considers mutual sequestration between anti-apoptotic and pro-apoptotic BCL-2 proteins, and between anti-apoptotic and BH3-only sensitizer/activator BCL-2 proteins, and direct activation of pro-apoptotic BCL-2 proteins by BH3-only proteins (Fig. 1). We construct a mathematical model based on a modified version of the ‘Embedded together’ framework, which pools sensitizer and activator BH3-only proteins. The model tracks the dynamics of three variables: the concentration of BH3-only proteins, *x*, anti-apoptotic BCL-2 proteins, *y*, and pro-apoptotic BCL-2 proteins, *z*.

**Figure 1.**
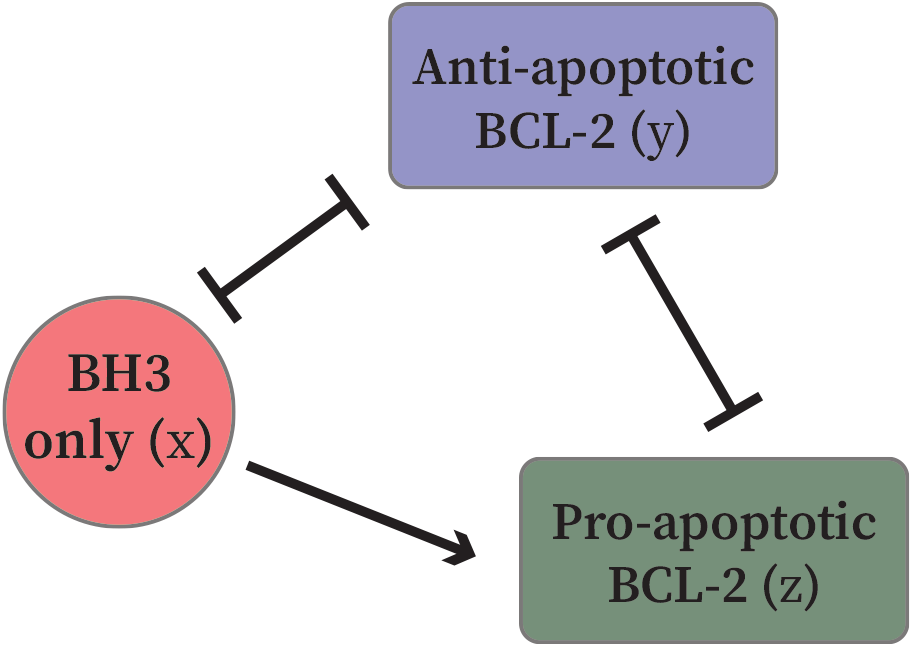
Schematic diagram of the modified ‘Embedded together’ model. Anti-apoptotic BCL-2 mutually inhibits pro-apoptotic and BH3-only BCL-2, while BH3-only BCL-2 activates pro-apoptotic BCL-2. Mutual inhibition/sequestration is represented by the blunt arrowhead and activation by the sharp arrowhead.

The deterministic system of equations governing BCL-2 protein interaction is given by:

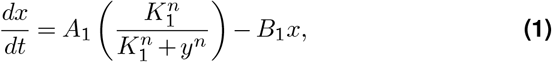

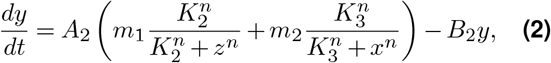

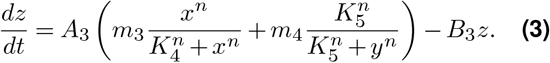

where the feedback between respective BCL-2 proteins, both inhibition and activation, is governed by a Hill function. This modeling choice is supported by experimental evidence showing that BH3-only proteins binding to BAX can induce conformational changes that enhance subsequent binding events, indicative of cooperative activation mechanisms Kale et al. (2018). Additionally, quantitative analyses of interactions between BCL-2 proteins and BH3 mimetics have reported steep, sigmoidal binding curves that could only be reproduced using cooperative models Osterlund et al. (2022).

The parameters *A*_*i*_ (*i* = 1, 2, 3) describe the maximum production rates of BH3-only proteins, anti-apoptotic BCL-2 proteins, and pro-apoptotic BCL-2 proteins, respectively. The parameters *K*_*i*_ (*i* = 1, …, 5) correspond to the dissociation constant, which describes the sensitivity to each respective regulator. The relative contribution of each feedback to the production of the protein is represented by *m*_*i*_ (*i* = 1, …, 4). Hence, in this model, the variables *x, y*, and *z* represent the unbound (free) concentrations of BH3-only, anti-apoptotic, and pro-apoptotic BCL-2 proteins, respectively. Finally, the Hill coefficient, *n*, describes the cooperativity of regulatory interactions. The parameter values of the model are given in Table 1. Since intrinsic apoptotic signals activate BH3-only proteins to trigger MOMP Delbridge et al. (2016); Radha and Raghavan (2017); Adams and Cory (2018); Singh et al. (2019), we consider *A*_1_ to be a natural bifurcation parameter to study the underlying dynamics and structure of the system. Thus, *A*_1_ serves both as a bifurcation parameter and as a proxy for the strength of the intrinsic apoptotic signal. The bifurcation structures throughout this work were computed using BifurcationKit.jl Veltz (2020). The behavior of the model for different rates of BH3-only protein production can be summarized in the bifurcation diagram shown in Fig. 2 and Table 2.

**Table 1.** System parameters.

| Parameter | Value | unit |
| --- | --- | --- |
| $A_2$ | 400 | conc/time |
| $A_3$ | 450 | conc/time |
| $B_i$ ($i = 1, 2, 3$) | 1.0 | 1/time |
| $m_i$ ($i = 1, 3$) | 1.0 | |
| $m_j$ ($j = 2, 4$) | 0.1 | |
| $K_i$ ($i = 1, \dots, 5$) | 1.0 | conc |
| $n$ | 2.0 | |

**Table 2.** Stability regimes as function of the bifurcation parameter, *A*_1_.

| $A_1$ range | Stable branches | Behavior |
| --- | --- | --- |
| (0, 4.958) | 1 | monostable |
| (4.958, 631.806) | 2 | bistable |
| (631.806, 2598.411) | 3 | tristable |
| (2598.411, 9872.774) | 2 | bistable |
| $> 9872.774$ | 1 | monostable |

**Figure 2.**
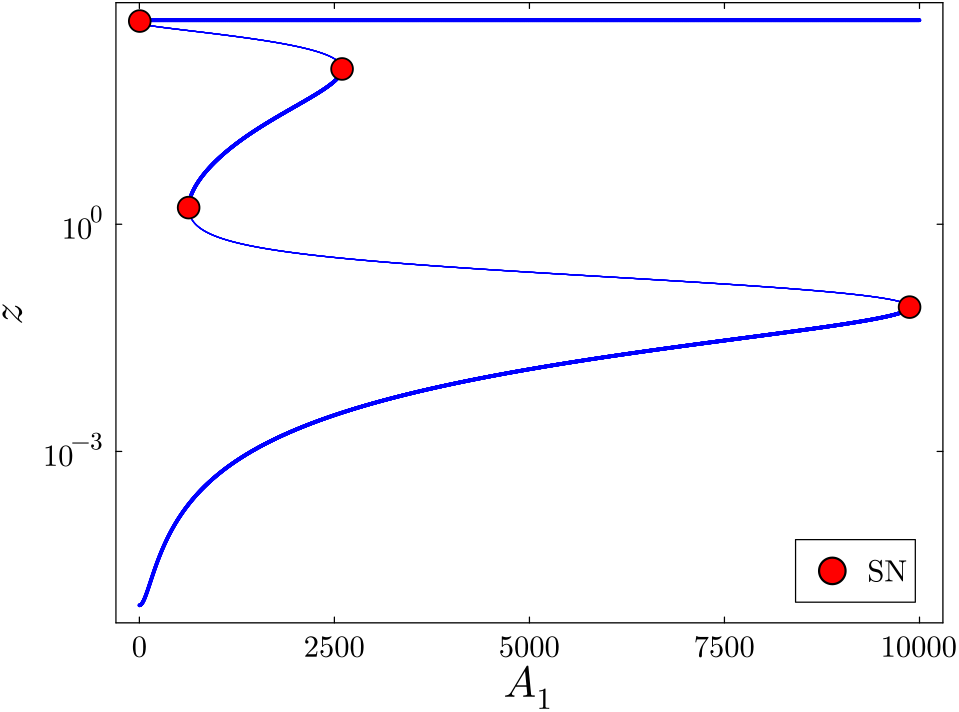
Partial bifurcation diagram of the model. The stable and unstable branches of the equilibrium solution as a function of *A*_1_ are shown as thick and thin curves, respectively. The response variable, *z*, is shown on a log scale. The red dots correspond to saddle-node bifurcations (SN) where the stability of the equilibrium solution changes.

The tristable region in our model captures cellular plasticity, wherein the same intrinsic apoptotic stimuli can drive cells toward three distinct stable steady states, namely, cell survival, senescence, or apoptosis, depending on the variability of initial conditions or intrinsic noise. In contrast, the bistable regions represent a binary switch, where cells have the choice between survival and apoptosis. Beyond these regimes, the system becomes monostable: for weak intrinsic stimuli (*A*_1_ *<* 4.958), cells uniformly survive, while for strong intrinsic stimuli (*A*_1_ *>* 9872.744), cells undergo apoptosis, nullifying internal variability. The transitions between various stable regions, and distinct cell fates, are marked saddle-node bifurcations (red dots, Fig. 2).

### Nondimensionalized model

To investigate how different stability regimes influence cell fate decisions, we analyze their robustness through parametric perturbations and phase space exploration. For enhanced biological relevance and numerical tractability, we derive a nondimensionalized version of the system.

We choose a reference concentration *K*_r_ to scale all concentration variables. Define:

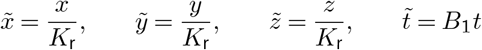

Substitute and define nondimensional parameters:

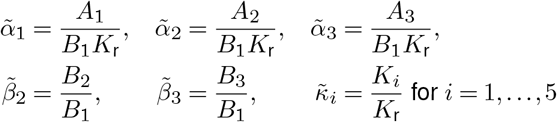

We choose *K*_r_ = 10 ensuring that all terms in the Hill functions remain 𝒪 (1) and keeping 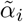 within a reasonable order of magnitude 𝒪 (103). Omitting the tildes for clarity, the nondimensionalized system is given by,

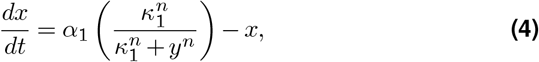

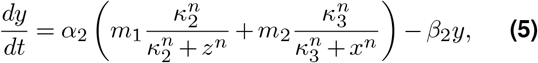

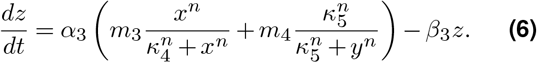

In the nondimensionalized system, the corresponding bifurcation parameter is *α*_1_ = *A*_1_*/*(*B*_1_*K*_r_). A partial bifurcation diagram is shown in Fig. S1

## Results

### BCL-2 protein expression control of multistability

BCL-2 proteins exhibit significant heterogeneous expression across different cell types, tissues, and pathological conditions. Consequently, this variability contributes to the differential treatment responses observed in therapies that target BCL-2 proteins Um (2015); Kale et al. (2018); Tessoulin et al. (2019); Smith et al. (2019); Qian et al. (2022); Perez-Serna et al. (2023). We investigate how the structural robustness and boundaries of the stability regimes change with the expression of BCL-2 protein family members. To this end, we perform a codimension-2 continuation tracking the saddle-node (SN) bifurcations in Fig. 2 in parameter space, revealing parameter regions that support tristable and bistable regimes.

Figure 3A shows changes in the stability regimes as *α*_1_ and *α*_2_ are varied, while Figs. 3B, C demonstrate stability regime changes as *α*_1_ and *α*_3_, and *α*_2_ and *α*_3_ are varied, respectively. Figure S2 displays these regions across a broader parameter range.

**Figure 3.**
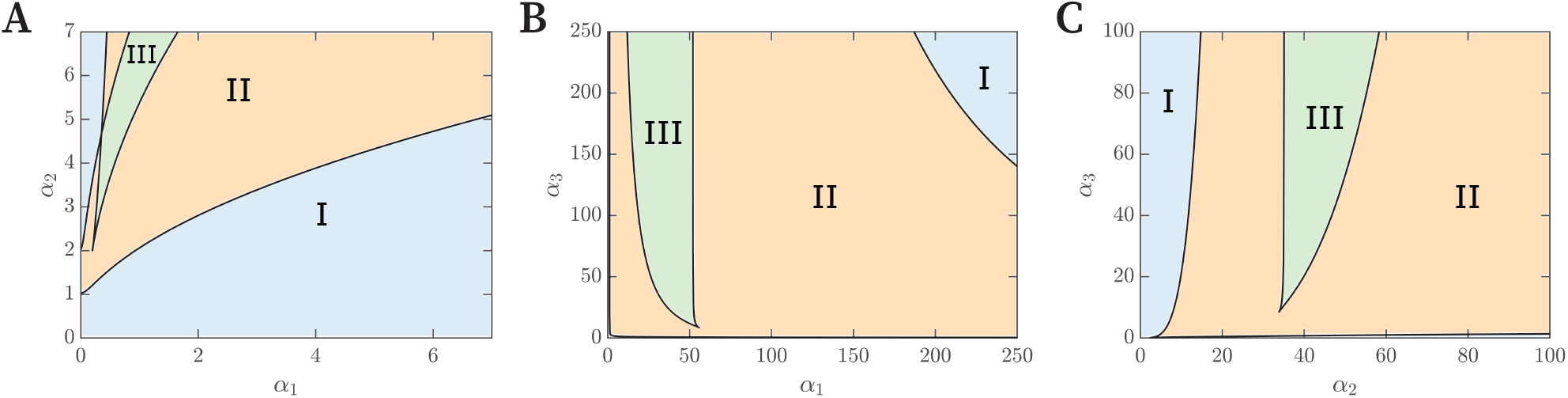
The bifurcation sets summarizes the changes in stability regimes as parameters corresponding to BCL-2 protein family expression (*α*_1_, *α*_2_ and *α*_3_) are varied. Regions where the system exhibits monostability are shown in blue and labeled I, regions of bistability are shown in orange and labeled II, while regions of tristability are shown in green and labeled III.

The bifurcation structure of the BCL-2 signaling model as defined by Eqs. (4)–(6) reveal a nuanced and asymmetric dependence on the key regulatory input, namely, BCL-2 expression. In (*α*_1_, *α*_2_)-space, bistability and tristability are robustly preserved across a broad range of parameter values. Multiple SN bifurcations persist in (*α*_1_, *α*_2_)-space, suggesting that cell fate decisions between survival, senescence, and apoptosis are strongly tunable by modifying the expression of BH3-only proteins and anti-apoptotic BCL-2 proteins. Both BH3-only and anti-apoptotic BCL-2 proteins are dynamically regulated by *p53* and under endoplasmic reticulum (ER) stress Hemann and Lowe (2006); Singh et al. (2019). In contrast, codimension-2 continuation in (*α*_1_, *α*_3_) and (*α*_2_, *α*_3_)-spaces demonstrates that multistable regimes are constrained to narrow ranges of *α*_3_ values, revealing a sharp threshold in pro-apoptotic BCL-2 proteins. Dynamically, this suggests a parameter-sensitive bifurcation geometry, where *α*_3_ serves as a bifurcation organizer, so that small changes in *α*_3_ can collapse multistable regimes. This property establishes parameter *α*_3_ (production of pro-apoptotic BCL-2 proteins) as a critical gatekeeper for apoptosis commitment. Biologically, this is consistent with the decisive role of pro-apoptotic BCL-2 proteins, BAX and BAK, whose release from anti-apoptotic BCL-2 or activation by BH3-only proteins induces irreversible MOMP. The observed parametric asymmetry in the multistable landscape illustrates how the network integrates continuous upstream signals (*α*_1_ and *α*_2_) with a threshold-like effector (*α*_3_) to control cell fate decisions with both sensitivity and robustness.

### Feedback sensitivity controls multistable robustness

In addition to protein abundance, cell fate is also governed by the affinity of BCL-2 proteins for binding partners Kale et al. (2018). Therefore, we systematically evaluate how the stability and transition between cell fate regimes are modulated by variations in BCL-2 affinities (dissociation constants; *κ*_*i*_, for *i* = 1, …, 5 in the model). Figure 4A suggests that multistable regimes are robustly maintained as BH3-only protein expression and their sensitivity to anti-apoptotic BCL-2 inhibition is varied. For *α*_1_ *>* 3, the tristable region persists for *κ*_1_ ≈ 0.1, while two regions of bistability are preserved for *κ*_1_ ≈ (0, 0.1) and *κ*_1_ ≈ (0.1, 1.0). Similarly, Fig. 4B displays the existence of multistable regions in (*α*_2_, *κ*_3_)-space. For an intermediate range of anti-apoptotic BCL-2 protein expression (*α*_2_ ≈ (12, 47)) tristable and bistable regimes persist for a wide range of sensitivity to BH3-only proteins. Interestingly, only when anti-apoptotic BCL-2 expression is low (*α*_2_ *<* 12) or high (*α*_2_ *>* 47), the system exhibits monostability, corresponding to cell survival or apoptosis.

**Figure 4.**
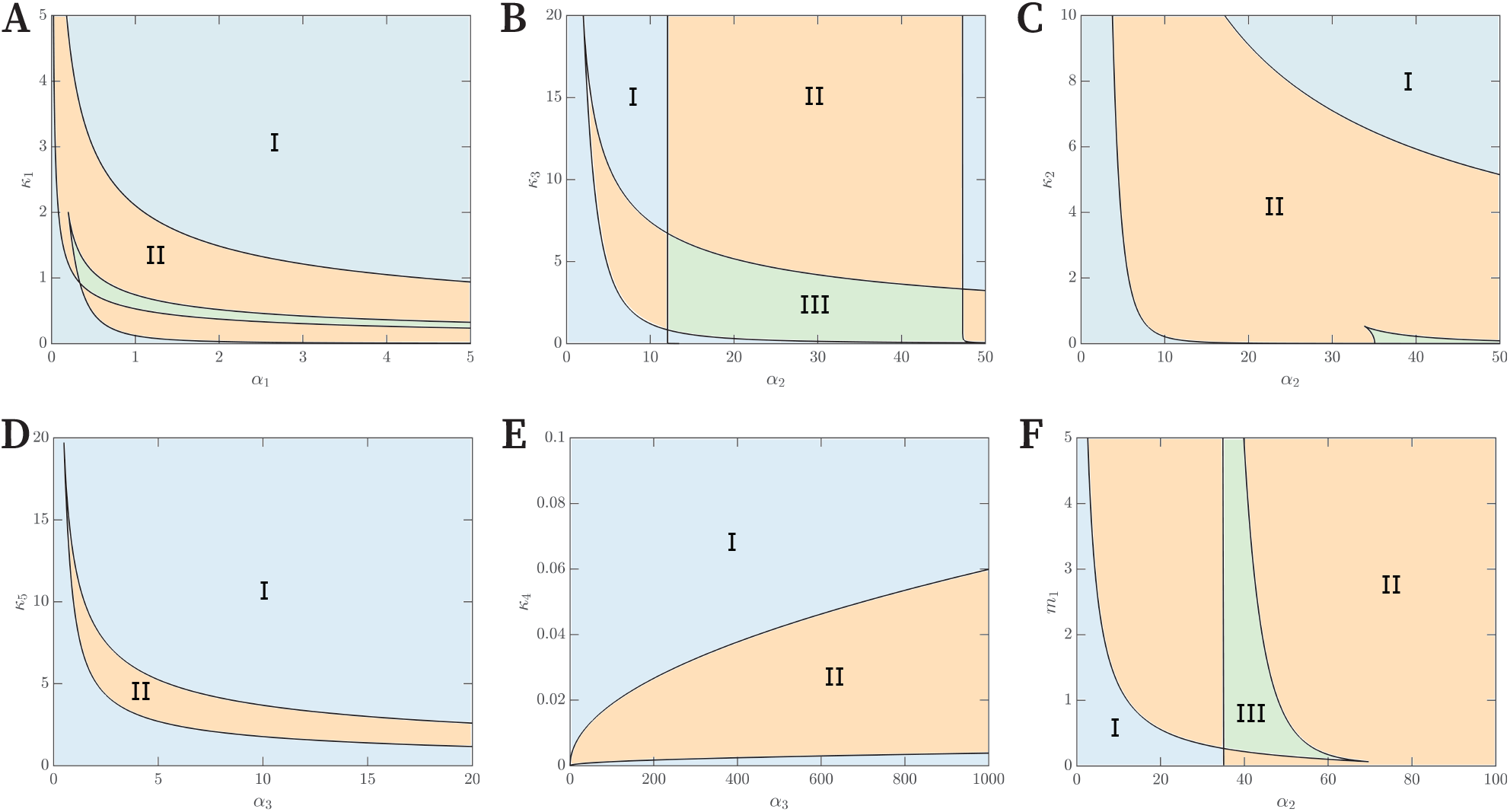
The bifurcation sets summarizes the changes in stability regimes as parameters corresponding to feedback sensitivity are varied. Regions where the system exhibits monostability are shown in blue and labeled I, regions of bistability are shown in orange and labeled II, while regions of tristability are shown in green and labeled III.

**Figure 5.**
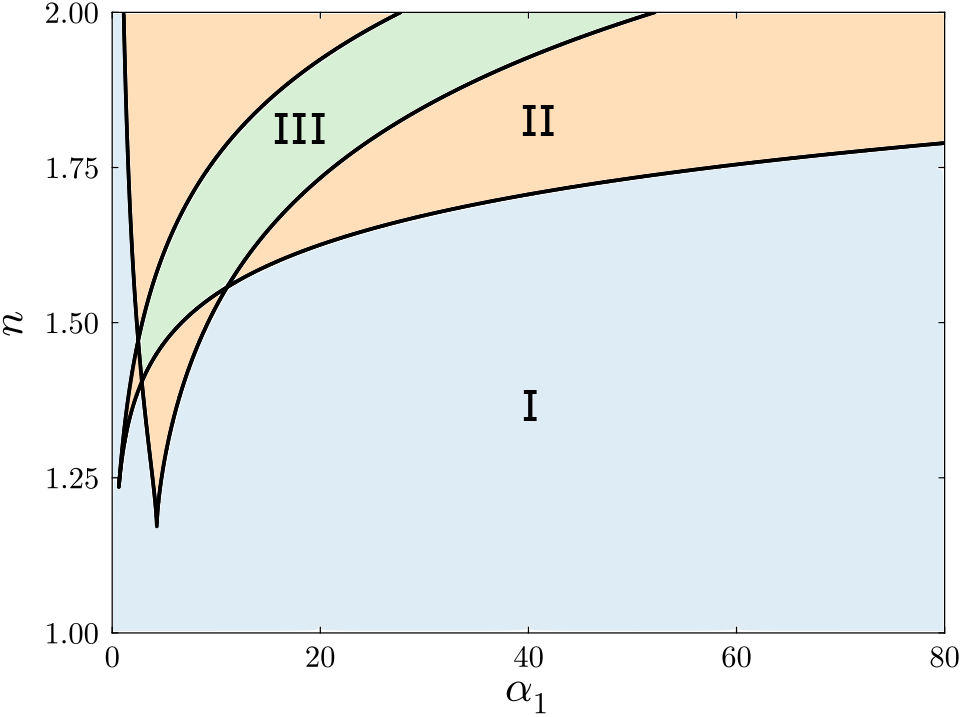
The bifurcation set summarizes the change in stability regimes in (*α*_1_, *n*)-space. Regions where the system exhibits monostability are shown in blue and labeled I, regions of bistability are shown in orange and labeled II, while regions of tristability are shown in green and labeled III.

Moreover, there are no tristable regimes for *κ*_1_, *κ*_3_ = 0, corresponding to absence of feedback between antiapoptotic BCL-2 proteins and BH3-only proteins and between BH3-only proteins and anti-apoptotic BCL-2 proteins, respectively. Therefore, for tristability to exist within the system, there has to be mutual sequestration of BH3-only and anti-apoptotic BCL-2 proteins, consistent with the ‘Embedded together’ modeling framework Shamas-Din et al. (2013).

Therapeutically, this suggests that in systems where anti-apoptotic BCL-2 proteins are overexpressed, leading to senescent or pro-survival states, BH3 mimetics with high anti-apoptotic BCL-2 affinity could force the system into apoptosis by driving the system across these bifurcation boundaries. These data provide further evidence that cell fate decisions can be modified by tweaking BH3-only protein and anti-apoptotic BCL-2 protein activation and expression, and their mutual sequestration.

In contrast, the tristable regime exists within a limited region for intermediate anti-apoptotic BCL-2 protein expression (35 *< α*_2_ *<* 60) and is finely tuned by proapoptotic BCL-2 feedback, *κ*_2_ (Fig 4C). The bistable regime, on the other hand, exists for a wide range of anti-apoptotic BCL-2 expression and sensitivity to pro-apoptotic BCL-2 inhibition. Figure 4D presents a structurally symmetric feedback, since anti-apoptotic BCL-2 and pro-apoptotic BCL-2 proteins mutually inhibit each other. The absence of a tristable regime reflects the simpler dynamics of pro-apoptotic BCL-2 signaling, demonstrating the threshold-driven all-or-nothing response of pro-apoptotic BCL-2 signaling. These findings underscore the therapeutic potential of targeting bifurcation thresholds to control cell fate decision-making. Figure 4E separates (*α*_3_, *κ*_4_)-space into monostable and bistable regimes. For low pro-apoptotic BCL-2 protein expressions (low *α*_3_) or low BH3-only feedback sensitivity (high *κ*_4_), the system remains in a monostable regime. However, when the BH3-only feedback becomes sufficiently strong, the system transitions to a bistable regime. This is consistent with the concept of apoptotic priming, where BH3-only proteins overcome a threshold, priming the cell for apoptosis King et al. (2022). Moreover, for *κ*_4_ = 0, the feedback from BH3only proteins to pro-apoptotic BCL-2 proteins is a constant, saturating at *m*_3_. Therefore, *m*_3_ acts as a constitutive activator of pro-apoptotic BCL-2 proteins, aligning with the ‘Direct activation’ framework, where BH3-only proteins (eg. BIM and NOXA) directly activate proapoptotic BCL-2 proteins (eg. BAX and BAK) ShamasDin et al. (2013).

The relative contribution of pro-apoptotic BCL-2 feedback on anti-apoptotic BCL-2 production is shown in Fig. 4F. Multistable regimes exist for large portions of the parameter space, highlighting the necessity of anti-apoptotic and pro-apoptotic BCL-2 protein competition for multistability. The ‘Unified’ BCL-2 framework posits that sequestration between anti-apoptotic BCL2 and pro-apoptotic BCL-2 proteins is more efficient in preventing apoptosis than sequestration between antiapoptotic BCL-2 and BH3-only proteins Llambi et al. (2011). In the model, the relative sequestration between anti-apoptotic BCL-2 proteins and BH3-only proteins is set to *m*_2_ = 0.1. The tristable region in Fig. 4F, emerges above *m*_1_ *>* 0.1, suggesting that phenotypic plasticity, where cells may undergo survival, senescence, or apoptosis, requires anti-apoptotic BCL-2 to preferentially interact with pro-apoptotic BCL-2 proteins over BH3-only proteins. However, for bistability regimes, this is not a requirement.

### Cooperativity control of multistable regimes

Exploring the bifurcation structure of our BCL-2 signaling network as a function of cooperativity reveals sharp transitions in the system’s dynamical landscape. For *n <* 1.22, the system remains monostable for all values of *α*_1_, indicating a regime where cooperativity is too weak to sustain multiple stable steady states. However, as *n* is increased beyond 1.22, the system acquires both bistable and tristable regimes, depending on the value of *α*_1_. Bistability arises at both low and high *α*_1_ values, while tristability is confined to intermediate ranges. For *n >* 1.5, these structures are more robust, with regions of multistability expanding and persisting for a wide range of parameter space. Thus, cooperative interactions in the regulatory feedback loops allow a high degree of dynamical plasticity and robustness to the network. Biologically, this aligns with the role of BH3-only proteins as a sensitizer. Whilst weak intrinsic apoptotic signal, characterized by low BH3-only protein activation, commits the cell to survival, a strong intrinsic apoptotic signal commits the cell to cell death. However, for intermediate BH3-only protein activation, the cell is ‘primed’ for but not committed to apoptosis, characterized by a multistable state.

Importantly, these results suggest that systems based purely on mass-action kinetics, without sequestration mechanisms or complex interaction topologies to introduce effective cooperativity, are unable to support multistability. Thus, incorporating Hill-type kinetics is not merely a modeling convenience but a reflection of underlying molecular mechanisms that shape the qualitative behavior of the system. This emphasizes the importance of cooperativity in biological regulation, especially in the context of senescence and apoptosis, for generating rich decision-making dynamics.

### Robustness of cell fate decision-making

We have performed an extensive analysis of the underlying bifurcation structure of our coarse-grained BCL2 signaling model, uncovering regions of monostability and multistability. While deterministic bifurcation analysis reveals the existence and stability of multiple attractors under parametric perturbations, it does not capture how intrinsic noise, ubiquitous in cellular signaling systems, influences cell fate decisions among these attractors.

By incorporating stochasticity into the dynamics via stochastic differential equations (SDEs), we investigate how intrinsic noise and initial conditions collectively shape cell fate decisions in regions of multistability. This allows us to map basins of attraction under stochastic dynamics and visualize how a unique parameter set can lead to cellular states such as survival, senescence, and apoptosis.

To model intrinsic noise, we reformulate the deterministic system as an SDE using the chemical Langevin approximation, where noise scales with the square root of the reaction fluxes Gillespie (2000). The general form of the SDE for species concentrations **u** = (*x, y, z*)*T* is,

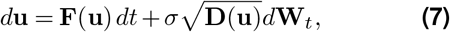

where **F**(**u**)= **F**+(**u**) − **F**− (**u**) represents the deterministic dynamics (Eqs. (4)–(6)), **D**(**u**) is a diagonal matrix with elements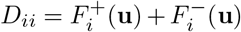, the sum of production and degradation rates for protein *i, σ* controls the noise amplitude and **W**_*t*_ is the standard Wiener process. Single-cell gene expression data reveals variations BCL-2 protein expression ranges between 5% and 40% Correia et al. (2015); Kueh et al. (2016); Qian et al. (2022). Here, simulations are performed with multiplicative noise introduced in the protein production and degradation terms, with *σ* = 0.05.

We perform stochastic simulations across a range of values of intrinsic apoptotic signal (*α*_1_). Each value of *α*_1_ corresponds to a distinct bifurcation regime in the deterministic system. For each regime, we simulate the SDE over a grid of initial conditions in relevant statespace projections, (*x*_0_, *y*_0_)-pairs with *z* = 0 fixed, running multiple replicates to account for the variability in the noise-induced dynamics. The final value of proapoptotic BCL-2 (*z*) is used to classify the cell’s fate into survival, senescence, or apoptosis. Model simulations were computed using DifferentialEquations.jl Rackauckas and Nie (2017).

Figure 6A, shows, for low BH3-only activation/expression (*α*_1_ = 0.5), all trajectories converge to low free pro-apoptotic BCL-2 expression, reflecting robust cell survival (colored red). In contrast, the tristable regime (Fig. 6B, *α*_1_ = 40) exhibits three distinct attractors corresponding to survival (red), senescence (pink) and apoptosis (blue), with basins of attraction delineated in state space. Thus, cells starting from different internal conditions or perturbed by intrinsic fluctuations may irreversibly commit to distinct cell fates, highlighting a plausible mechanism for cellular heterogeneity.

**Figure 6.**
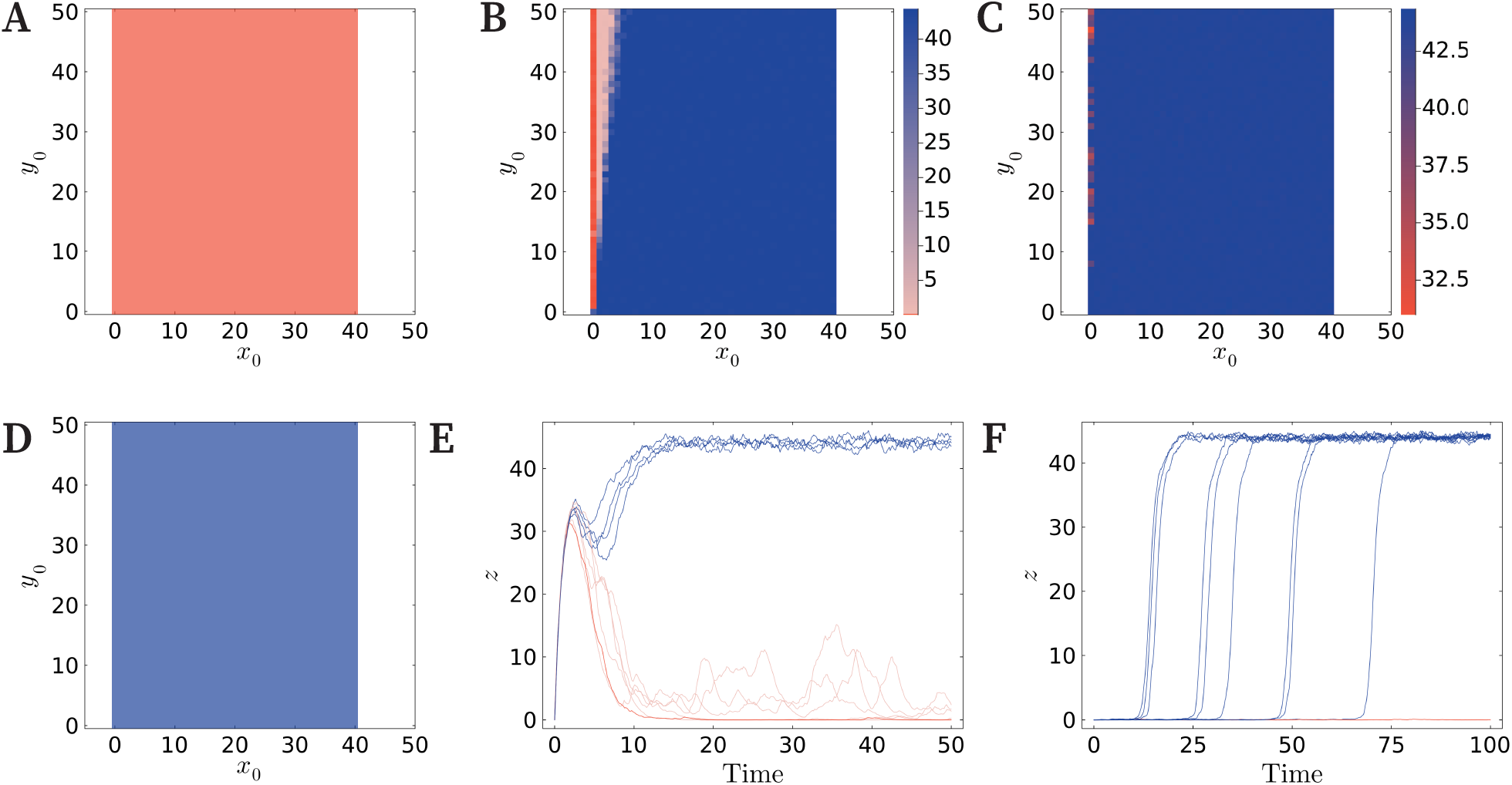
Cell fate decisions under intrinsic noise and varying initial conditions. Within regions of multistability identified in Fig. 2 we perform stochastic simulations, recording basins of attractions for the final pro-apoptotic BCL-2 state (*z*) under varying initial BH3-only and anti-apoptotic BCL-2 concentrations ((*x*_0_, *y*_0_)-pairs). For all simulations *z*_0_ = 0. (A) For *α*_1_ low, simulations suggest cells are restricted to a survival state regardless of initial condition. In the tristable and bistable regions (intermediate *α*_1_), (B) and (C), respectively, cells may undergo survival, senescence or apoptosis, depending on the initial protein concentrations. For *α*_1_ high, cells are restricted to an apoptosis state regardless of initial conditions. (E) In the tristable regime, time series simulations for (*x*_0_, *y*_0_)= (1, 4) show some cells undergoing survival or senescence, while the majority of cells undergo apoptosis. Similarly, in the bistable regime (F), the majority of cells undergo apoptosis while some undergo survival for ((*x*_0_,*y*_0_) = (0, 33). Red color corresponds to cell survival, pink to senescence and blue to apoptosis. For simulations, the coefficient of variation is 5%. 10 replicates are simulated for each initial condition pair.

Similarly, the bistable regime (Fig. 6C, *α*_1_ = 500), constrains cellular decision-making to survival or apoptosis depending on initial BH3-only and anti-apoptotic BCL2 protein concentrations. Survival tends to dominate for low initial BH3-only levels (*x*_0_), whereas apoptosis becomes the predominant outcome as *x*_0_ increases. Figure 6D demonstrates that high BH3-only activation/expression (*α*_1_ = 1100) enforces irreversible commitment to apoptosis, regardless of initial BH3-only and anti-apoptotic BCL-2 proteins concentrations.

To further illustrate how intrinsic noise can drive divergent cell fates from the same initial molecular state, we performed stochastic time series simulation in both the tristable and bistable regimes. For each regime, we selected a single initial condition and ran ten replicates of the stochastic system. Figure 6E, demonstrate how trajectories starting from the same initial condition can lead cells to survival (red), senescence (pink) or apoptosis (blue), with individual realizations committing to different attractors over time. Similarly, in the bistable regime (Fig. 6F), trajectories diverge toward either cell survival or apoptosis. Notably, our model simulations capture the variable time-to-switch behavior observed experimentally Albeck et al. (2008). Collectively, these results emphasize how multistability and noise can encode probabilistic cell fate decisions in cell populations and offer a mechanistic basis for non-genetic heterogeneity in cell fate decision-making. These findings underscore how cell fate decisions in multistable systems are shaped by intrinsic molecular noise.

### Mapping to mechanistic models

Next, we map the coarse-grained bifurcation structure of our phenomenological BCL-2 model onto mechanistic models of BCL-2 signaling that explicitly model biochemical interactions. This approach enables us to interpret the dynamical regimes as emergent regimes grounded in specific molecular architectures, bridging systems-level insights with molecular detail. While previous mechanistic models of apoptotic signaling networks have used bifurcation analyses, these models have typically only revealed bistability within their networks Legewie et al. (2006); Chen et al. (2007); Cui et al. (2008); Zhao et al. (2015). In contrast, our model reveals a much richer dynamical landscape, including robust tristable regimes. By analyzing the bifurcation structure of other mechanistic models, with a particular focus on BCL-2 signaling, and positioning them within this broader dynamical framework, we may identify which molecular features are necessary or sufficient to reproduce complex decision-making dynamics. This cross-scale mapping not only enhances the biological interpretability of our coarse-grained model but also enables the classification of mechanistic models by the qualitative behaviors they support.

Our analysis specifically examines the BCL-2 signaling components within established models of the apoptotic network. Cui *et al*. utilize mass-action kinetics to model interactions between BCL-2 family proteins, proposing a hierarchy of models with increasing complexity Cui et al. (2008). Their most comprehensive framework (termed the ‘Direct Action II’ model) dynamically tracks concentrations of pro-apoptotic and antiapoptotic BCL-2 proteins, as well as BH3-only sensitizer and activator proteins. This model incorporates: (i) sequestration between BCL-2 protein families, (ii) competitive displacement from heterodimers, and (ii) BAX autoactivation (Fig. 7). The model equations of the Cui *et al*. model are shown in the supplement. Comparative analyses revealed that this formulation exhibits the most robust bistability among their proposed modeling schemes, suggesting its utility for capturing apoptotic threshold behaviors.

**Figure 7.**
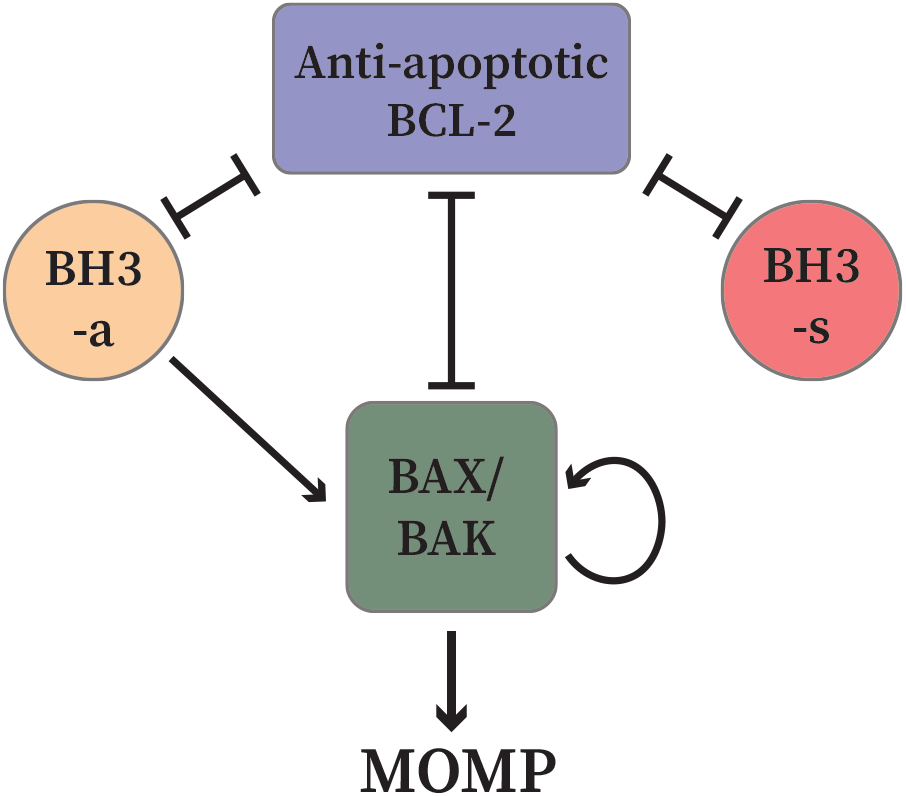
Schematic diagrams of the Cui *et al*. model Cui et al. (2008). The model accounts for dynamic changes in the concentrations of pro-apoptotic (green) and anti-apoptotic (blue) BCL-2 and BH3-only activator (orange) and sensitizer (red) proteins. Mutual inhibition/sequestration is represented by blunt arrowheads and activation by sharp arrowheads.

While bistability in this system has been well characterized, the potential for tristability in this system remains underexplored. Understanding how the kinetic parameters govern multistability and robustness is crucial for elucidating cell fate decisions. Therefore, we systematically perturb key parameters in the Cui *et al*. model to assess their influence on the hysteresis range and the emergence of tristability. The partial bifurcation diagram for the unperturbed Cui *et al*. model is shown in Fig. S3, black curve.

We first examined parameters controlling BH3-only activator (BH3a) and anti-apoptotic BCL-2 (BCL-2) interactions. Parameters *k*_4_ and *k*_7_ determine the sequestration and displacement of pro-apoptotic BAX, respectively. Increasing both *k*_4_, BH3a-BCL-2 binding, and *k*_7_, BH3a-mediated BAX displacement, synergistically widened the hysteresis range, enhancing the system’s resistance to noise (Fig. S3, blue dotted curve). However, these modifications were unable to introduce tristability, suggesting that BH3a-driven feedback alone is insufficient to establish a third stable state.

We next investigated BAX activation dynamics, where *k*_15_ governs autoactivation and *k*_16_ regulates BAX oligomerization. Interestingly, reducing *k*_16_ increased hysteresis, but eliminating it destroyed bistability, revealing that weak but nonzero oligomerization is essential for maintaining a bistable switch. This implies that cooperative BAX aggregation creates a threshold for irreversible apoptosis commitment, but its absence renders the system monostable. The simultaneous modification of *k*_15_ and *k*_16_ revealed unexpected dynamics in the system’s bistable regime, widening the hysteresis range but inducing a counterintuitive response in the upper stable branch, where increased BH3a production (*p*_2_) yields a decrease in MOMP level (Fig. S3, blue dashed curve). This non-monotonic behavior may arise from competitive inhibition, where high BH3a levels might sequester BCL-2 more effectively, paradoxically limiting BAX from oligomerizing or saturation effects in BAX activation, where excessive BH3a leads to rapid but incomplete oligomerization. Furthermore, this inverse relationship may also arise from a timescale separation between BAX activation and oligomerization becoming more pronounced at extreme *p*_2_ values. These results suggests the existence of an optimal BH3a concentration for maximal MOMP commitment.

We then tested the role of BH3 sensitizer (BH3s) proteins which competitively inhibit BCL-2. Increasing *k*_13_, BH3s-mediated BAX displacement, slightly expanded hysteresis but failed to introduce tristability, indicating that BH3s primarily buffers BCL-2 rather than establishing new stable states (Fig. S3, blue solid curve). BH3s binding to BCL-2 (*k*_9_) had negligible effects on the hysteresis range (*results not shown*), underscoring the importance of BH3a-driven regulation in this system. Similarly, perturbations of BCL-2 production (*p*_3_) did not confer significant robustness to the bistability of the system (*results not shown*).

Finally, we modified five parameters (*k*_4_, *k*_7_, *k*_13_, *k*_15_ and *k*_16_) simultaneously (Fig. S3, green curve). To introduce tristability into the system, we attempt to create two bistable switches, a lower switch (survival to senescence) controlled by parameters *k*_4_ and *k*_7_ and an upper switch (senescence to apoptosis) controlled by parameters *k*_13_, *k*_15_ and *k*_16_. Modifying these parameters collectively widened the hysteresis range, enhancing the robustness of bistability in the system. However, these perturbations failed to induce tristability, indicating that while competitive BH3-only binding and BAX activation kinetics sharpen the apoptotic switch, additional regulatory layers are required to establish a third stable state. This suggests that robust cell-fate decisions in apoptosis rely on tightly coupled BH3-only driven priming and pro-apoptotic activation, whereas a metastable intermediate (senescent) phenotype may require independent control mechanisms.

To bridge the gap between phenomenological abstraction and mechanistic modeling, we hybridize our phenomenological model by incorporating mechanistic terms derived from mass-action kinetics. This approach retains the analytical tractability and interpretability of the phenomenological framework while embedding biochemically grounded interactions where the mechanistic detail is well-understood. In particular, we replace Hill-type regulation in Eqs (4)-(6) with MichaelisMenten-type terms by setting the Hill coefficient to one in pathways where the molecular binding or sequestration is thought to proceed via non-cooperative interactions. This hybridization provides a more faithful representation of molecular control mechanisms without reintroducing the full complexity of the mechanistic model, thereby facilitating the exploration of multistability in a biologically-informed, but computationally efficient modeling framework.

In the first hybridization, we replace Hill-type regulation with Michaelis-Menten kinetics for interactions between BH3-only proteins and anti-apoptotic BCL-2 proteins, as well as between anti-apoptotic BCL-2 and pro-apoptotic BCL-2 proteins, reflecting non-cooperative binding dynamics. Only the BH3-only-dependent activation of pro-apoptotic BCL-2 proteins retained phenomenological Hill-type regulation. This hybrid model generated bistability, but the hysteresis range was extremely narrow (*α*_1_ ≈ 1.047 to *α*_1_ ≈ 1.072), indicating that noncooperative sequestration alone is insufficient to produce a robust bistable switch (Fig. 8A). This suggests that simple, non-cooperative mutual inhibition between BH3-only and anti-apoptotic BCL-2 proteins, while sufficient to create a theoretical survival-to-apoptosis switch, does not recapitulate the strong irreversibility observed in physiological apoptosis. The narrow hysteresis implies that cells operating under such a regime would be highly sensitive to fluctuations in BH3-only signaling, potentially leading to stochastic, rather than decisive, cell-fate transitions.

**Figure 8.**
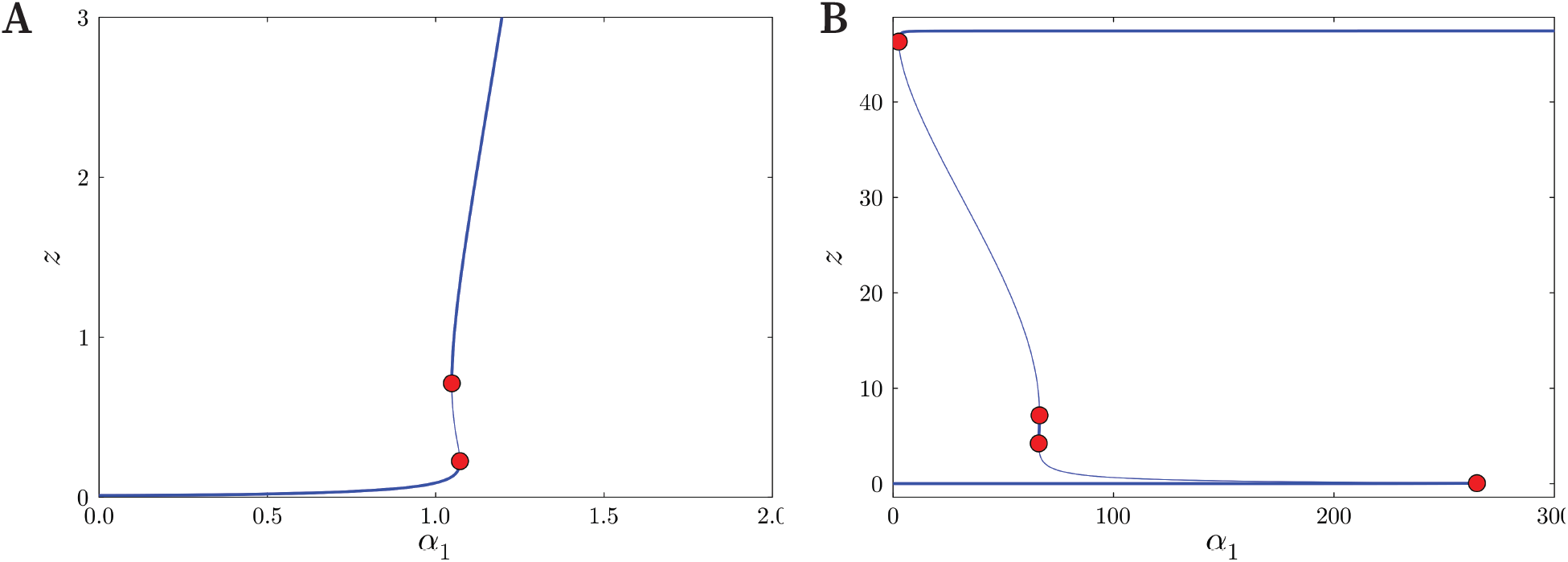
Partial bifurcation diagrams of a hybridized models. (A) Partial bifurcation diagram of the hybridized model, with interactions between BH3-only and antiapoptotic BCL-2, and between anti-apoptotic BCL-2 and pro-apoptotic BCL-2 proteins modeled with Michaelis-Menten-type equations (*n* = 1). The system exhibits bistability as the bifurcation parameter, *α*_1_, is varied. The system demonstrates a sharp, sensitive switch between the survival (lower stable branch) and apoptosis (upper stable branch with a narrow hysteresis range. (B) Partial bifurcation diagram of the hybridized model, with only interactions between anti-apoptotic BCL-2 and pro-apoptotic BCL-2 proteins modeled with Michaelis-Menten-type equations. This system exhibits tristability for a narrow range of the bifurcation parameter, *α*_1_. The stable and unstable branches of the equilibrium solution as a function of *α*_1_ are shown as thick and thin curves, respectively. The red dots correspond to saddle-node bifurcations where the stability of the equilibrium solution changes.

In the second hybridization, we introduced cooperativity for BH3-only and anti-apoptotic BCL-2 interactions, reflecting experimental evidence that BH3-only proteins exhibit cooperative binding to BCL-2 Osterlund et al. (2022). Interactions between anti-apoptotic and proapoptotic BCL-2 proteins remain non-cooperative. This modification generated tristability, with three distinct stable branches (Fig. 8B). However, the intermediate branch, within the tristable regime, was extremely narrow (*α*_1_ ≈ 66.06 to *α*_1_ ≈ 66.41), suggesting that while BH3-only and anti-apoptotic BCL-2 interactions enable multistability, the tristable state is metastable and highly sensitive to parameter perturbations. This suggests that cells can enter a transiently senescent phenotype, where partial anti-apoptotic BCL-2 inhibition delays proapoptotic BCL-2 activation, but the system remains predisposed to apoptosis upon minor additional stress.

These hybrid models demonstrate that cooperativity between BH3-only and anti-apoptotic BCL-2 proteins is critical for generating multistability, while noncooperative anti-apoptotic and pro-apoptotic BCL-2 interaction limits the robustness of the tristable regime. This suggests that physiological tristability likely requires additional layers of control to expand the stability range of the senescent phenotype.

## Discussion

Apoptosis and senescence are critical processes in development, aging, and disease, with dysregulation contributing to pathologies such as cancer, neurodegenerative disorders, and cardiovascular diseases Kerr et al. (1972); Elmore (2007); Hayflick (1965); Childs et al. (2014). While apoptosis results in cell death, senescence leads to irreversible cell cycle arrest, with the choice between these fates often determined by stress levels Chen et al. (2000); Song et al. (2005); Vousden and Lane (2007); Saleh et al. (2020). The BCL-2 protein family, key regulators of apoptosis, also modulates senescence, highlighting the interconnectedness of these pathways Roger et al. (2021); Basu (2022).

The BCL-2 family includes pro-apoptotic (e.g., BAX, BAK), anti-apoptotic (e.g., BCL-2, BCL-xL), and BH3only proteins, which regulate mitochondrial outer membrane permeabilization (MOMP) to control apoptosis Hata et al. (2015); Kalkavan and Green (2018); Delbridge et al. (2016). Anti-apoptotic BCL-2 proteins inhibit MOMP, while BH3-only proteins counteract this inhibition, promoting apoptosis. Interestingly, BCL-2 upregulation has been linked to senescence, while, in some contexts, senescent cells exhibit resistance to apoptosis despite reduced BCL-2 levels Wang (1995); Bladier et al. (1997); Tombor et al. (2003). This suggests that BCL-2 proteins play a dual role in determining cell fate.

Senescent cells often overexpress anti-apoptotic BCL-2 members, contributing to their survival and implicating these proteins in age-related diseases and therapy resistance Childs et al. (2014); Zhu et al. (2016); Schmitt et al. (2022). Therapy-induced senescence, for example, may promote tumor progression, underscoring the need for targeted interventions against BCL-2 proteins in cancer and senescence-related conditions Baar et al. (2017); Wang et al. (2022). Thus, BCL-2 proteins represent promising therapeutic targets for modulating apoptosis and senescence in disease contexts.

Despite progress in understanding apoptosis and senescence, the mechanisms governing their cell fate decisions, particularly the regulatory role of BCL-2 proteins, remain unclear. While computational models have explored BCL-2-mediated apoptosis signaling and, to a lesser extent, senescence pathways, existing frameworks fail to integrate both processes or capture the tristable decisions between survival, senescence and apoptosis Fussenegger et al. (2000); Eissing et al. (2004); Chong et al. (2018). To address this gap, we develop a novel BCL-2-dependent model that systematically analyzes the dynamic interactions governing these fate choices, with a focus on how BCL-2 family proteins confer robustness to cellular decision-making. This approach provides a more comprehensive framework to dissect the molecular switches between apoptosis, senescence, and survival.

Given the significant heterogeneity in BCL-2 protein expression across cell types and disease states, influencing therapeutic responses Um (2015); Kale et al. (2018); Tessoulin et al. (2019), we analyzed how varying BCL-2 family member expression alters system robustness and stability regimes. Our codimension-2 bifurcation analysis (Fig. 2) reveals broad tristable and bistable regions in (*α*_1_, *α*_2_)-space (Figs. 3A, B), demonstrating tunable cell fate decisions through BH3-only and antiapoptotic BCL-2 regulation. In contrast, multistability collapses sharply with small changes in pro-apoptotic BCL-2 levels (*α*_3_), establishing it as a critical threshold for apoptosis commitment. This parametric asymmetry highlights how the network combines graded upstream signals (e.g., p53-or ER stress-regulated *α*_1_/*α*_2_) with a switch-like *α*_3_ response to ensure robust yet sensitive fate determination. These results align with BAX/BAK’s decisive role in MOMP and explain how variable BCL-2 expression may differentially prime cells for apoptosis, senescence, or survival.

Beyond protein expression levels, our analysis reveals that cell fate decisions are critically dependent on binding affinities between BCL-2 family members. The system maintains robust multistability when mutual sequestration exists between BH3-only and anti-apoptotic BCL-2 proteins (Figs. 4A,B), with tristability requiring intermediate anti-apoptotic BCL-2 expression and bidirectional feedback. This aligns with the ‘Embedded to-gether’ framework Shamas-Din et al. (2013), emphasizing the necessity of competitive interactions for phenotypic plasticity. Therapeutically, these findings suggest BH3 mimetics could overcome anti-apoptotic BCL-2 overexpression by shifting systems across bifurcation boundaries into apoptosis.

In contrast, pro-apoptotic BCL-2 regulation exhibits threshold-driven bistability (Figs. 4C-E), where BH3-only proteins must exceed a critical threshold to trigger apoptosis, consistent with apoptotic priming King et al. (2022). Notably, tristability emerges only when antiapoptotic BCL-2 preferentially sequesters pro-apoptotic BCL-2 proteins over BH3-only proteins (Fig. 4F), supporting the ‘Unified’ framework’s prediction that such interactions dominate apoptotic blockade Llambi et al. (2011). These results collectively demonstrate how binding affinities and feedback architectures encode distinct regulatory logics, graded control for survival/senescence versus switch-like apoptosis commitment, providing a roadmap for targeted interventions.

Our analysis of cooperativity in the BCL-2 network reveals that multistability emerges only when *n >* 1.22, with robust bistable and tristable regimes forming for *n >* 1.5. The absence of multistability for *n <* 1.22 underscores that cooperativity, whether from sequestration, feedback loops, or Hill-type kinetics, is essential for generating the plasticity observed in cell fate decisions, rather than being a mere modeling artifact. These dynamics align with BH3-only proteins’ role as sensitizers, where graded signals are converted into discrete fate choices through cooperative molecular interactions.

This mirrors biological decision-making: weak BH3-only signals favor survival, strong signals trigger apoptosis, and intermediate signals create a ‘primed’ multistable state.

To link our deterministic analysis with biological reality, we incorporated stochasticity through chemical Langevin equations, revealing how noise and initial conditions jointly govern fate decisions in multistable regimes. In tristable regions, identical cells diverge into survival, senescence, or apoptosis based on stochastic fluctuations and initial anti-apoptotic BCL-2 and BH3-only protein concentrations (Fig. 6B), while bistable systems exhibit probabilistic switching between survival and apoptosis (Fig. 6C). Notably, time-course simulations demonstrate that single initial conditions can yield divergent fates across replicates (Figs. 6E,F). These results establish that intrinsic noise, when coupled with multistable dynamics, generates non-genetic heterogeneity, illustrating how genetically identical cells under uniform stimuli can adopt distinct fates. The model thus provides a quantitative framework linking molecular fluctuations to population-level phenotypic diversity in apoptosis-senescence decisions.

Our study bridges phenomenological and mechanistic modeling approaches to dissect how BCL-2 network architecture governs cell fate decisions. While established mechanistic models robustly capture bistability through BH3-only and anti-apoptotic BCL-2 sequestration and BAX autoactivation, our analysis reveals their inability to generate tristability through parameter variation alone, highlighting the need for additional regulatory layers to establish metastable senescent states. Hybrid models incorporating cooperative BH3-only and anti-apoptotic BCL-2 interactions successfully induced tristability, but with an extremely narrow intermediate regime, suggesting physiological systems likely employ complementary mechanisms: feedback loops, spatial compartmentalization of anti-apoptotic and pro-apoptotic BCL-2 proteins or post-translational modifications, to stabilize senescence.

These findings collectively demonstrate that cell fate decisions emerge from an interplay of molecular specificity, nonlinear dynamics (cooperativity and hysteresis), and stochasticity. While BH3-only and anti-apoptotic BCL-2 cooperativity enables multistability, robust tristability requires further biological constraints. Our integrated framework not only deciphers how molecular interactions encode cell fate plasticity but also provides a blueprint for manipulating these decisions therapeutically. Ultimately, the BCL-2 network exemplifies how biological systems exploit dynamical complexity to translate noisy molecular signals into decisive, yet adaptable, cellular outcomes.

## Acknowledgments

IC is supported by the María de Maeztu Postdoctoral Fellowship. We thank the CERCA Program/Generalitat de Catalunya for institutional support. This research has been funded by the grant PID2021-127896OB-I00 funded by MCIN/AEI/10.13039/501100011033 ‘ERDF A way of making Europe’. This has been supported by the Spanish Research Agency (AEI), through the Severo Ochoa and Maria de Maeztu Program for Centres and Units of Excellence in R&D (CEX2020-001084-M).

## Data Availability

The simulations carried out in this study were performed using Julia computing language Bezanson et al. (2017). Bifurcation analyses were performed using Julia’s bifurcationkit.jl package Veltz (2020), while stochastic simulations were performed using Julia’s DifferentialEquations.jl package Rackauckas and Nie (2017). The Julia code to perform the bifurcation analyses and the stochastic simulations is available on GitHub under the Apache License 2.0 (https://www.github.com/Ielyaas/BCL-2-model).

