## Supplementary for "Dynamical analysis of a model of BCL-2-dependent cellular decision making"

### Supplementary Information

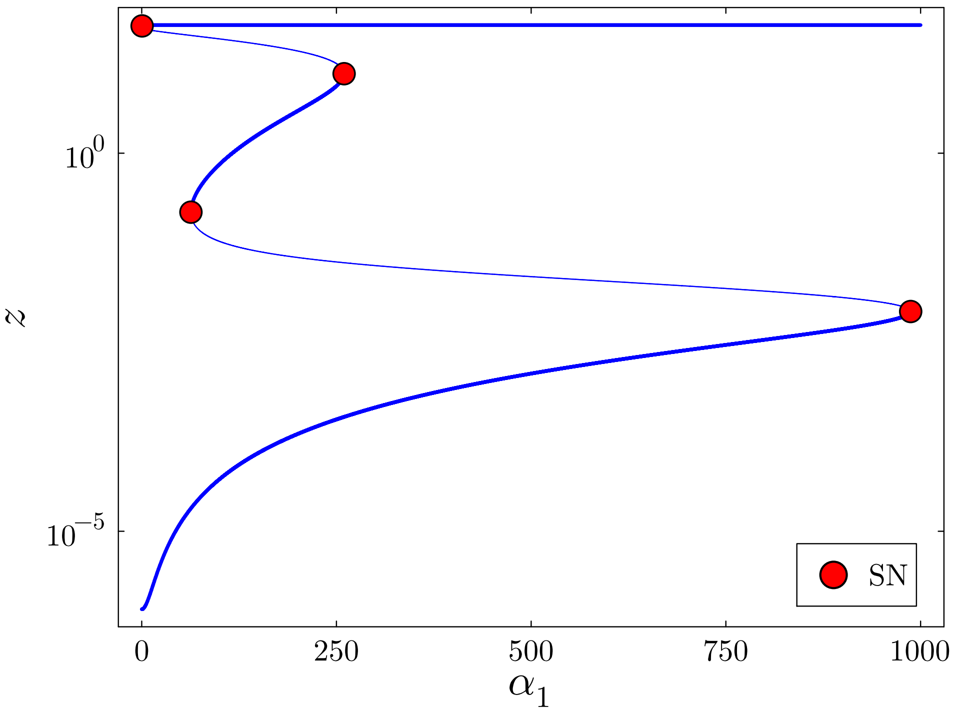


**Figure S1.** Partial bifurcation diagram of the nondimensionalized model. The stable and unstable branches of the equilibrium solution as a function of $\alpha_1$ are shown as thick and thin curves, respectively. The response variable, $z$, is shown on a log scale. The red dots correspond to saddle-node bifurcations (SN) where the stability of the equilibrium solution changes.

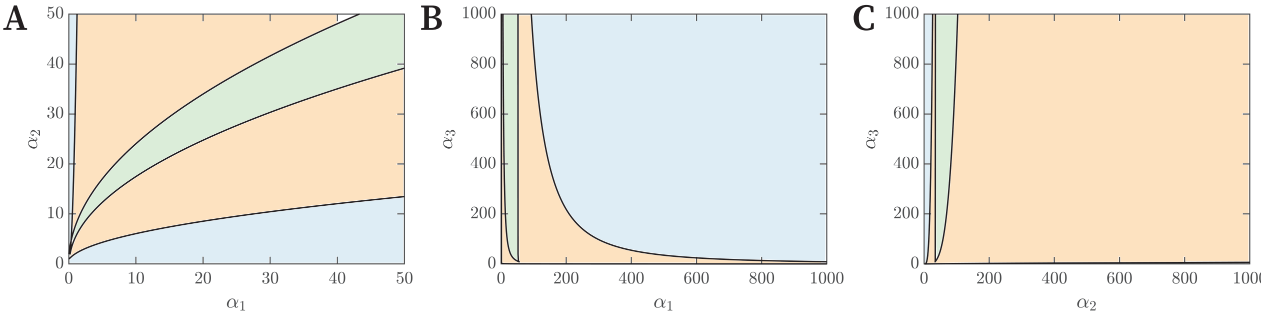


**Figure S2.** The bifurcation set summarizes the changes in stability regimes as BCL-2 expression parameters ($\alpha_1$, $\alpha_2$ and $\alpha_3$) are varied. The blue, orange and green regions correspond to monostable, bistable and tristable regimes, respectively.

**Cui Model**

The system of equations of the Cui *et al.* model is given by

$$\begin{aligned}
\frac{d[\mathrm{BAX}]}{dt} &= p_1 - u_1[\mathrm{BAX}] - k_1[\mathrm{BAX}][\mathrm{BH3a}] - k_{15}[\mathrm{BAX}][\mathrm{aBAX}],\\
\frac{d[\mathrm{aBAX}]}{dt} &= -u_2[\mathrm{aBAX}] + k_1[\mathrm{BAX}][\mathrm{BH3a}] - k_2[\mathrm{aBAX}][\mathrm{BCL2}] + k_3[\mathrm{aBAX_BCL2}]\\
&\quad - k_6[\mathrm{aBAX}][\mathrm{BH3a_BCL2}] + k_7[\mathrm{aBAX_BCL2}][\mathrm{BH3a}] - k_8[\mathrm{aBAX}]\\
&\quad + k_{13}[\mathrm{BH3s}][\mathrm{aBAX_BCL2}] - k_{14}[\mathrm{aBAX}][\mathrm{BH3s_BCL2}]\\
&\quad - k_{15}[\mathrm{BAX}][\mathrm{aBAX}] - 2k_{16}[\mathrm{aBAX}]^2 + 2k_{17}[\mathrm{MOMP}],\\
\frac{d[\mathrm{BCL2}]}{dt} &= p_3 - u_4[\mathrm{BCL2}] - k_2[\mathrm{aBAX}][\mathrm{BCL2}] + k_3[\mathrm{aBAX_BCL2}]\\
&\quad - k_4[\mathrm{BH3a}][\mathrm{BCL2}] + k_5[\mathrm{BH3a_BCL2}]\\
&\quad - k_9[\mathrm{BH3s}][\mathrm{BCL2}] + k_{10}[\mathrm{BH3s_BCL2}],\\
\frac{d[\mathrm{BH3a}]}{dt} &= p_2 - u_3[\mathrm{BH3a}] - k_1[\mathrm{BAX}][\mathrm{BH3a}] - k_4[\mathrm{BH3a}][\mathrm{BCL2}] + k_5[\mathrm{BH3a_BCL2}]\\
&\quad + k_6[\mathrm{aBAX}][\mathrm{BH3a_BCL2}] - k_7[\mathrm{aBAX_BCL2}][\mathrm{BH3a}]\\
&\quad + k_{11}[\mathrm{BH3s}][\mathrm{BH3a_BCL2}] - k_{12}[\mathrm{BH3a}][\mathrm{BH3s_BCL2}],\\
\frac{d[\mathrm{BH3a_BCL2}]}{dt} &= -u_5[\mathrm{BH3a_BCL2}] + k_4[\mathrm{BH3a}][\mathrm{BCL2}] - k_5[\mathrm{BH3a_BCL2}]\\
&\quad - k_6[\mathrm{aBAX}][\mathrm{BH3a_BCL2}] + k_7[\mathrm{aBAX_BCL2}][\mathrm{BH3a}]\\
&\quad - k_{11}[\mathrm{BH3s}][\mathrm{BH3a_BCL2}] + k_{12}[\mathrm{BH3a}][\mathrm{BH3s_BCL2}],\\
\frac{d[\mathrm{aBAX_BCL2}]}{dt} &= -u_6[\mathrm{aBAX_BCL2}] + k_2[\mathrm{aBAX}][\mathrm{BCL2}] - k_3[\mathrm{aBAX_BCL2}]\\
&\quad + k_6[\mathrm{aBAX}][\mathrm{BH3a_BCL2}] - k_7[\mathrm{aBAX_BCL2}][\mathrm{BH3a}]\\
&\quad - k_{13}[\mathrm{BH3s}][\mathrm{aBAX_BCL2}] + k_{14}[\mathrm{aBAX}][\mathrm{BH3s_BCL2}],\\
\frac{d[\mathrm{BH3s}]}{dt} &= p_4 - u_7[\mathrm{BH3s}] - k_9[\mathrm{BH3s}][\mathrm{BCL2}] + k_{10}[\mathrm{BH3s_BCL2}]\\
&\quad - k_{11}[\mathrm{BH3s}][\mathrm{BH3a_BCL2}] + k_{12}[\mathrm{BH3a}][\mathrm{BH3s_BCL2}]\\
&\quad - k_{13}[\mathrm{BH3s}][\mathrm{aBAX_BCL2}] + k_{14}[\mathrm{aBAX}][\mathrm{BH3s_BCL2}],\\
\frac{d[\mathrm{BH3s_BCL2}]}{dt} &= -u_8[\mathrm{BH3s_BCL2}] + k_9[\mathrm{BH3s}][\mathrm{BCL2}] - k_{10}[\mathrm{BH3s_BCL2}]\\
&\quad + k_{11}[\mathrm{BH3s}][\mathrm{BH3a_BCL2}] - k_{12}[\mathrm{BH3a}][\mathrm{BH3s_BCL2}]\\
&\quad + k_{13}[\mathrm{BH3s}][\mathrm{aBAX_BCL2}] - k_{14}[\mathrm{aBAX}][\mathrm{BH3s_BCL2}],\\
\frac{d[\mathrm{MOMP}]}{dt} &= -u_9[\mathrm{MOMP}] + k_{15}[\mathrm{BAX}][\mathrm{aBAX}] + k_{16}[\mathrm{aBAX}]^2 - k_{17}[\mathrm{MOMP}].
\end{aligned}$$

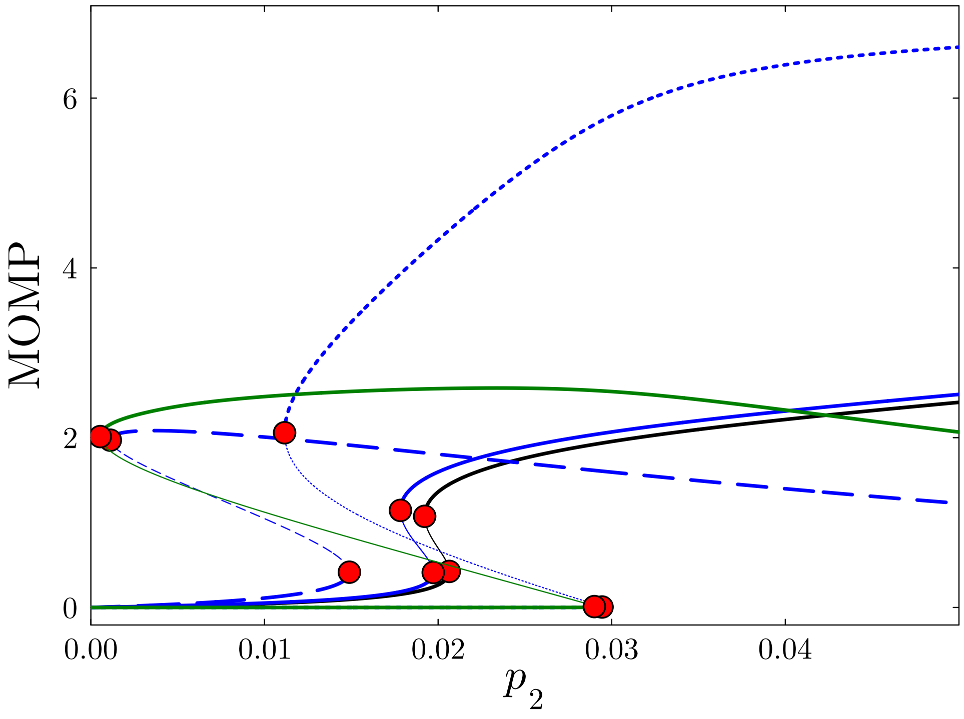


**Figure S3.** Robustness of bistability in the Cui *et al.* model Cui et al. (2008). Partial bifurcation diagrams are shown under different parametric perturbations. The black curve shows the bifurcation diagram of the base model. We assess the robustness of bistability by perturbing critical parameters controlling positive feedback loops. The solid blue curve corresponds to the bifurcation diagram when $k_{13}$, corresponding to BH3 sensitizer displacement of activated BAX, is modified. The dashed blue curve corresponds to the bifurcation diagram when parameters $k_{15}$ and $k_{16}$, corresponding to BAX activation and oligomerization are modified. The dotted blue curve corresponds to the bifurcation diagram when parameters $k_4$ and $k_7$, corresponding to competitive BH3-only - BCL-2 binding is modified. Finally, the solid green curve represents the bifurcation diagram when all five parameters ($k_4$, $k_7$, $k_{13}$, $k_{15}$ and $k_{16}$) are modified. With each modification we notice an increase in the hysteresis width as function of the bifurcation parameter $p_2$. The red dots correspond to saddle-node bifurcations (SN) where the stability of the equilibrium solution changes.
